# Genetic Explication of Impaired Insulin Resistance in Genesis of Metabolic Diseases

**DOI:** 10.1101/2024.05.02.592139

**Authors:** Naveed Iqbal Soomro, Syeda Marriam Bakhtiar

## Abstract

This study aims to investigate the role of impaired insulin resistance in the onset and progression of metabolic diseases such as prediabetes, diabetes, and cardiovascular diseases. Insulin resistance occurs when insulin is unable to effectively stimulate glucose uptake, and if the body is unable to produce sufficient insulin to compensate, type 2 diabetes may develop. This research endeavors to elucidate the molecular and genetic underpinnings of insulin resistance and its association with metabolic disorders. Employing various tools and databases, gene interaction data was procured through GeneMania, and pathway validation was conducted using KEGG. Construction of gene regulatory networks employed GEPHI 0.9.2, with centralities statistical analysis identifying hub genes. Enrichment analysis and literature validation substantiated the significance of these hubs, resulting in the refinement of the initially identified seven genes to five with interaction data. The implicated hub genes were discerned to play roles in inflammation, either directly or indirectly. Future prospects involve further genetic analysis across diverse populations, utilizing PCR to discern the allelic variations of these identified hub genes. Ultimately, this research may shed light on the underlying genetic and molecular mechanisms of insulin resistance and metabolic syndrome, and contribute to the development of targeted treatments for these conditions.

## Introduction

Metabolic diseases are a group of disorders that affect the body’s ability to process and use nutrients (such as carbohydrates, proteins, and fats) for energy and other essential functions (1). These disorders can be caused by genetic mutations, environmental factors, or a combination of both. Some examples of metabolic diseases include prediabetes that leads to diabetes type 2, obesity, hypertension, and cardiovascular diseases (CVDs) (2).

Prediabetes refers to a medical condition in which blood sugar levels are elevated, yet not to the point of being classified as type 2 diabetes (3). It acts as a warning signal that an individual might develop type 2 diabetes, a chronic disease with severe complications like kidney and heart damage (4–6). Prediabetes is a condition affecting approximately one in three adults in the United States and one in five adults worldwide, with higher prevalence rates observed in low- and middle-income countries (7).

Diabetes is a chronic metabolic disease associated with insulin abnormalities due to inadequate insulin secretion or ineffective use of insulin. According to the International Diabetes Federation (https://diabetesatlas.org/). 537 million adults are living with diabetes; Diabetes is responsible for 6.7 million deaths in 2021(7).

Obesity is a condition where there is an excessive accumulation of body fat, which is typically defined by having a BMI of 30 or higher (8). The prevalence of obesity has been steadily increasing worldwide, with over 650 million adults being classified as obese in 2016. This is a significant public health concern due to the increased risk of chronic diseases associated with obesity (9).

Hypertension, also known as high blood pressure, is a condition where the force of blood against the walls of arteries is persistently high, leading to a strain on the heart and an increased risk of health problems. It is estimated that 1.13 billion people worldwide had hypertension in 2015, and this number is expected to rise to 1.56 billion by 2025 (10).

Cardiovascular diseases (CVDs) are a collection of medical conditions that impact the heart and blood vessels (11). They are responsible for a significant proportion of global deaths, accounting for around one-third of all fatalities each year. Cardiovascular diseases (CVDs) are the leading cause of death globally, responsible for 32% of all deaths in 2019 (12).

These metabolic diseases may lead to severe conditions such as Metabolic syndrome (MetS). Metabolic syndrome is a cluster of conditions that increase the risk of developing cardiovascular diseases, type 2 diabetes, and other health problems. A diagnosis is typically made when a person has at least three of the following conditions: elevated fasting glucose, elevated blood pressure, elevated triglycerides, reduced HDL cholesterol, and abdominal obesity (13). The global prevalence of metabolic syndrome is estimated to be around 25%, with higher rates in urban areas and among older adults (14).

Insulin resistance plays a central role in the development of metabolic syndrome. Insulin is a hormone produced by the pancreas that helps to regulate the amount of glucose (sugar) in the blood. Insulin resistance occurs when the body’s cells become resistant to the effects of insulin, leading to high levels of glucose in the blood (15).

When insulin resistance develops, the pancreas responds by producing more insulin to overcome the resistance. This leads to higher-than-normal levels of insulin in the blood, a condition known as hyperinsulinemia. Over time, the pancreas may become exhausted and unable to produce enough insulin, leading to high blood sugar levels and eventually Pre-diabetes leading to diabetes. Insulin resistance also affects the body’s ability to use and store energy from food (6). This can lead to the accumulation of fat in the liver and other organs, which can contribute to the development of non-alcoholic fatty liver disease. Additionally, insulin resistance can increase inflammation in the body, which can contribute to the development of other health problems such as cardiovascular disease (16).

Insulin resistance and metabolic syndrome are influenced by a complex interplay of genetic and environmental factors. Various genes and genetic variations have been linked to these conditions, including PPARG, IRS1, IRS2, ADIPOQ, TNF, FTO, SIRT1, and GCKR (17–21). These genes are involved in various processes related to energy metabolism, adipocyte differentiation, lipid metabolism, and inflammation, among others (Figure 1). However, the genetic basis of insulin resistance and metabolic syndrome is not fully understood, and additional research is needed to unravel their complex genetic and environmental components.

**Figure 1:**
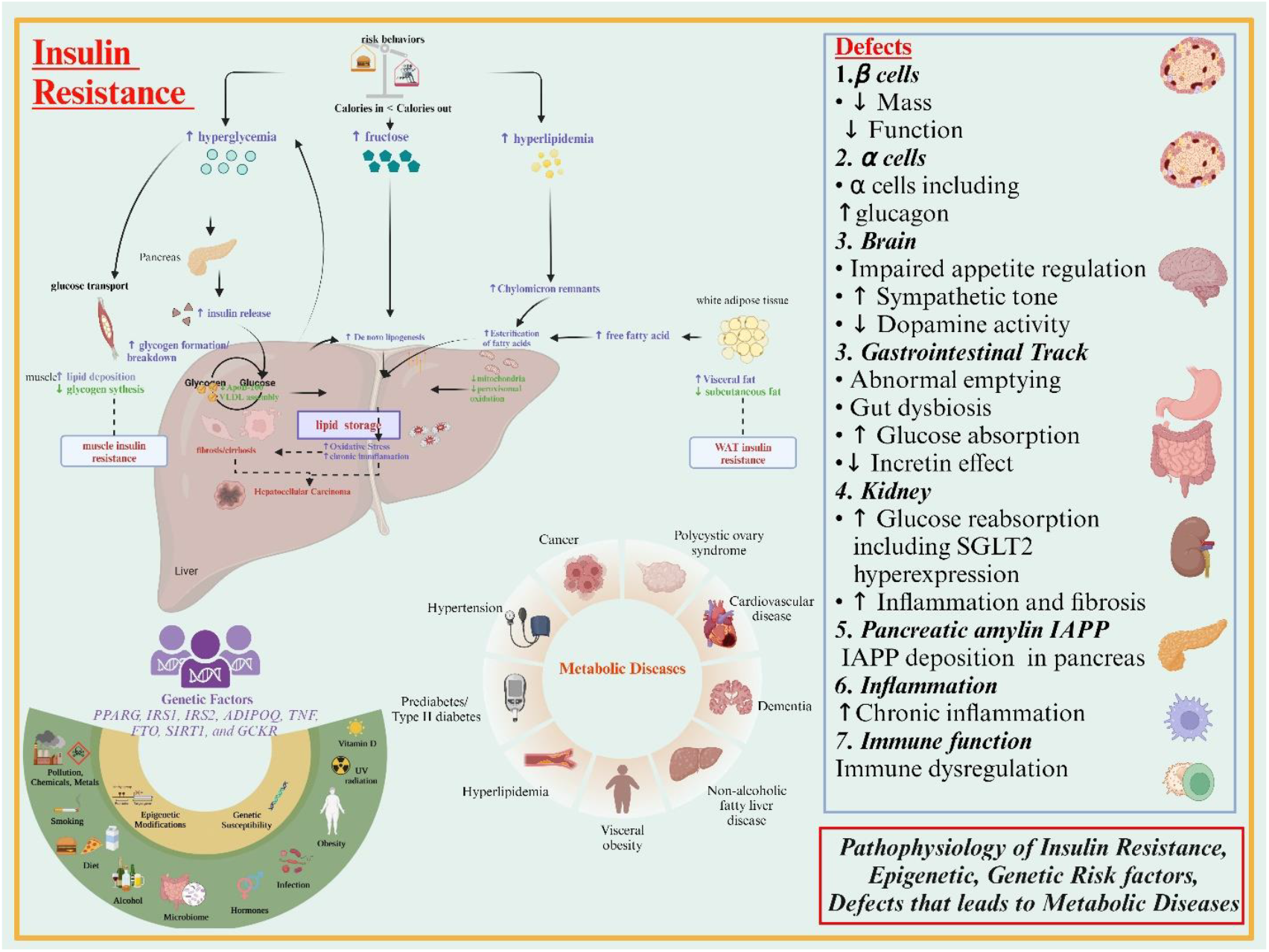
Role of Insulin Resistance in the prognosis of metabolic diseases

## Methodology

### Data collection

The top-down bioinformatics approach was used, core genes that are associated with insulin resistance and metabolic syndrome were primarily retrieved from literature i.e., PubMed (https://pubmed.ncbi.nlm.nih.gov/) as well as from pathways that are available in databases such as KEGG (https://www.genome.jp/kegg/pathway.html).

### Gene Interaction Networks

Gene interaction network with other genes and the function of genes were predicted through the online tool GeneMANIA (http://genemania.org/) (22). Nodes and edges of interacted genes were further processed in Excel to remove duplication.

### GGI network construction and identification of hub genes

The Excel file consisting of nodes and edges was uploaded to GEPHI 0.9.2 (https://gephi.org/) (23) and gene regulatory networks and interaction data were retrieved to find hub genes associated with insulin resistance. Data obtained from GEPHI was analyzed for values of closeness centrality (24), harmonic centrality (25), betweenness centrality (26), and eigncentrality were compared by adjusting threshold values to identify hub genes of every network.

### Functional annotation and pathway enrichment analysis

Enrichment analysis of hub genes was performed through the ToppGene tool (https://toppgene.cchmc.org/) (27). Hub genes were uploaded to the ToppGene tool for pathway enrichment based on gene ontology (GO) for biological processes, molecular function, and cellular components (28).

### Validation of hub genes

Finally, hub genes identified were further validated through previously published literature to investigate whether these genes are involved in inflammation directly or through any metabolic disease that causes inflammation. The overview of methodology implemented is presented in (Figure 2).

**Figure 2:**
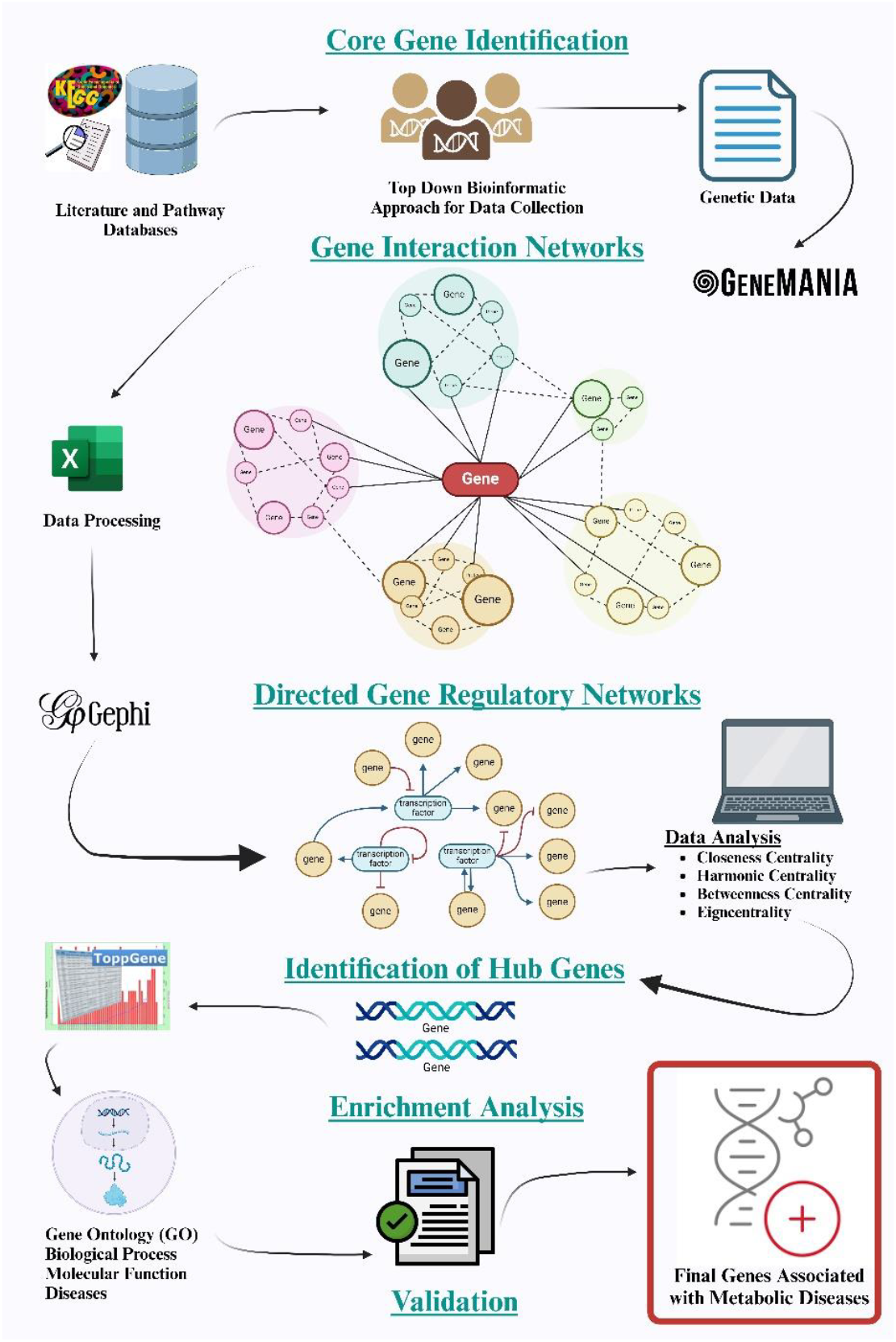
Methodology overview

## Results

### Identification of core genes

Seven core genes of insulin resistance were identified from literature and pathways of several organs such as adipose tissue, liver, and skeletal muscles. These genes are IRS1, IRS2 AKT2 functional in adipose tissue, liver, muscles, and heart, FOXO1 functions in the liver and heart, TNF-α, DAG, and IKK-β functions in the kidney, liver, and muscles (Table 1). The selection of core genes associated with metabolic diseases involved a thorough process. We began with a comprehensive literature review to identify consistently reported genes and prioritized those from Genome-Wide Association Studies (GWAS) with statistically significant associations. Our focus on functional relevance considered genes actively participating in metabolism and insulin signaling pathways. Transcriptomic studies and protein-protein interaction networks provided valuable insights. The rigorous evaluation included genetic validity through replication and consideration of clinical relevance. Meta-analyses and consensus among studies strengthened our selection. Notably, we prioritized genes experimentally validated through functional assays and animal studies, ensuring the identification of robust core genes associated with metabolic diseases.

**Table 1:** Core genes selected from published literature and pathways

| Sr No. | Genes | Organ |
| --- | --- | --- |
| 1 | IRS1 | Adipose tissue, liver, muscles, heart |
| 2 | AKT2 | Adipose tissue, liver, muscles, heart |
| 3 | FOXO1 | Liver, heart |
| 4 | TNF-α | Kidney, liver, muscle |
| 5 | DAG | Liver, muscle, kidney |
| 6 | IKK-β | Liver, muscle, kidney |
| 7 | IRS2 | Adipose tissue, liver, muscles, heart |

### sInteraction networks

After conducting a thorough literature and pathway analysis to verify the genes, we identified interactions among the genes using GeneMania (Figure 3) and downloaded the resulting interaction data for genes (supplementary data). While we were able to obtain interaction data for five of the seven genes, TNF-α and IKK-β genes were not recognized by GeneMania. In addition, we investigated networks that involved multiple pathways and retrieved the relevant pathways from KEGG (Kyoto Encyclopedia of Genes and Genomes) (Table 2).

**Table 2:**
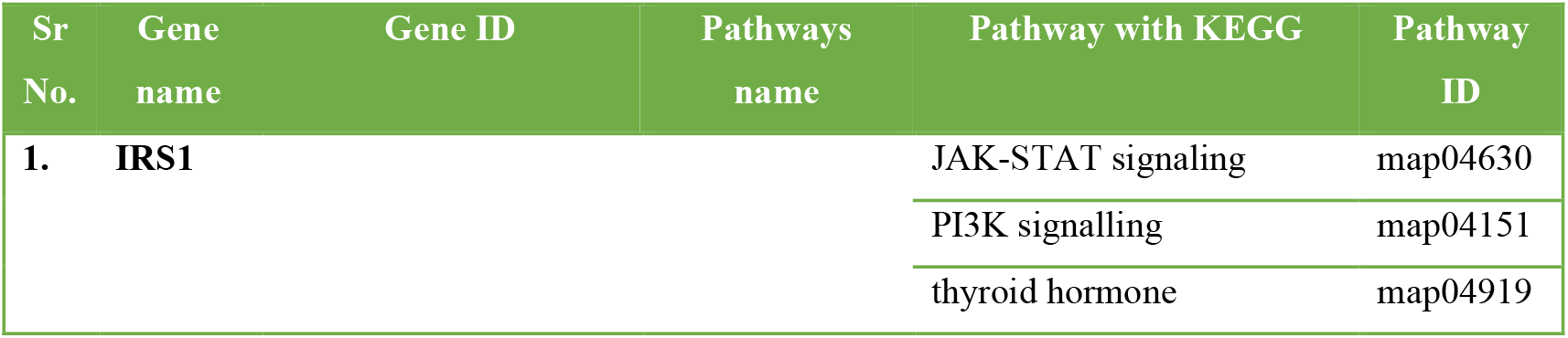
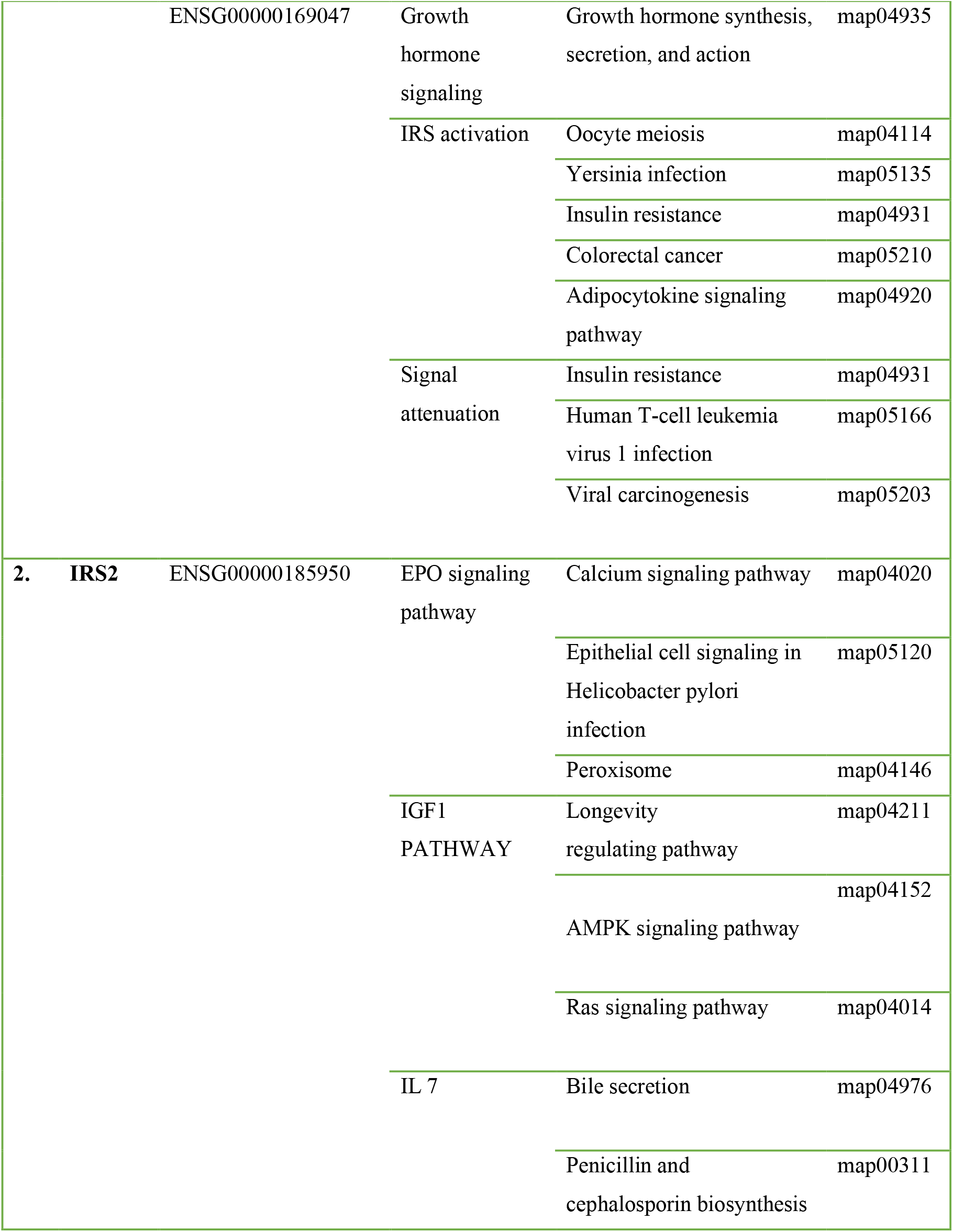
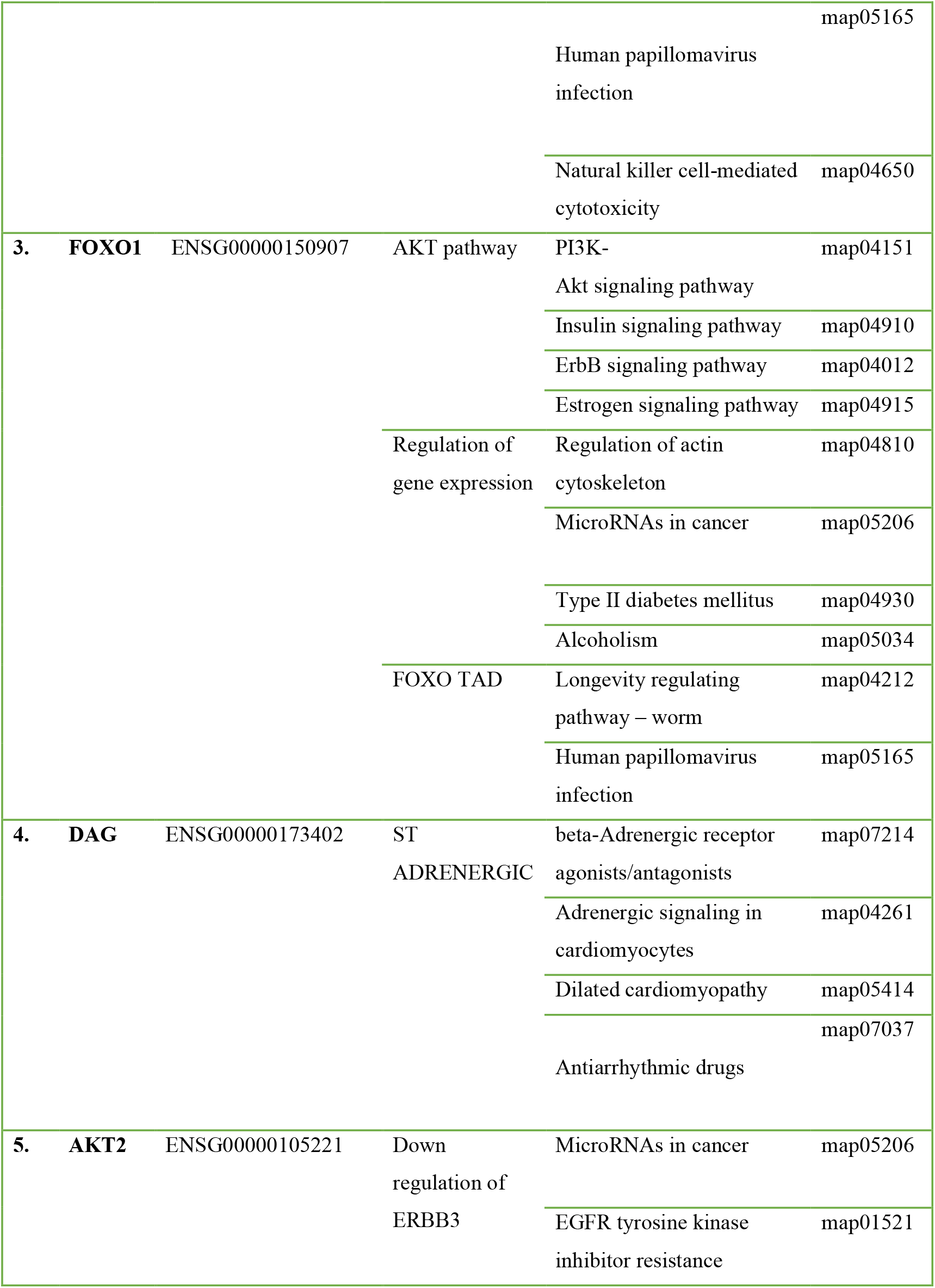
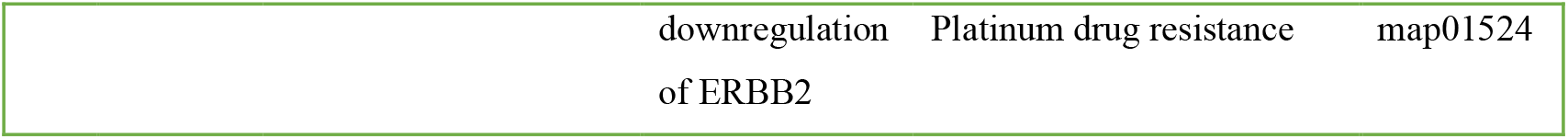
Pathways associated with genes.

**Figure 3:**
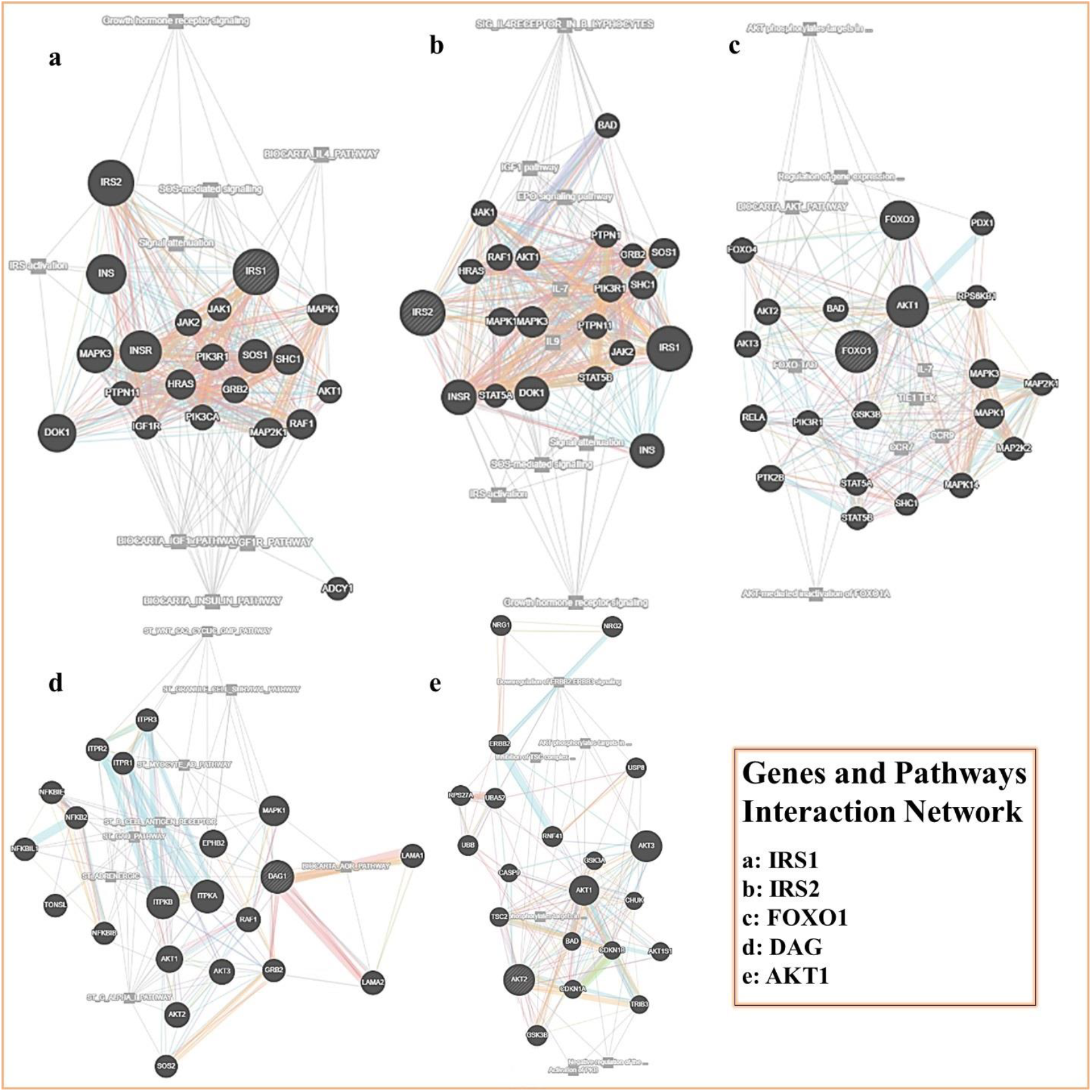
Genes and pathway interaction networks

### Identification of hub genes

We utilized a tool called Gephi 0.9.2 to develop and analyze Directed Gene Regulatory networks (Figure 4). We applied various statistical tests, including degrees and average weighted degrees, to determine the number of interactions between the genes. The centralities of the nodes in the graph were also evaluated, such as betweenness, which indicates the significance of a node in connecting different parts of the network, closeness, which measures the shortest distance between nodes, and harmonic, which assesses the influence of neighboring nodes. Through this analysis, we aimed to identify hub genes in the network, which are nodes with the highest centralities. These hub genes may play crucial roles in regulating the activity of other genes and pathways within the network, making them potential targets for further investigation and therapeutic intervention.

**Figure 4:**
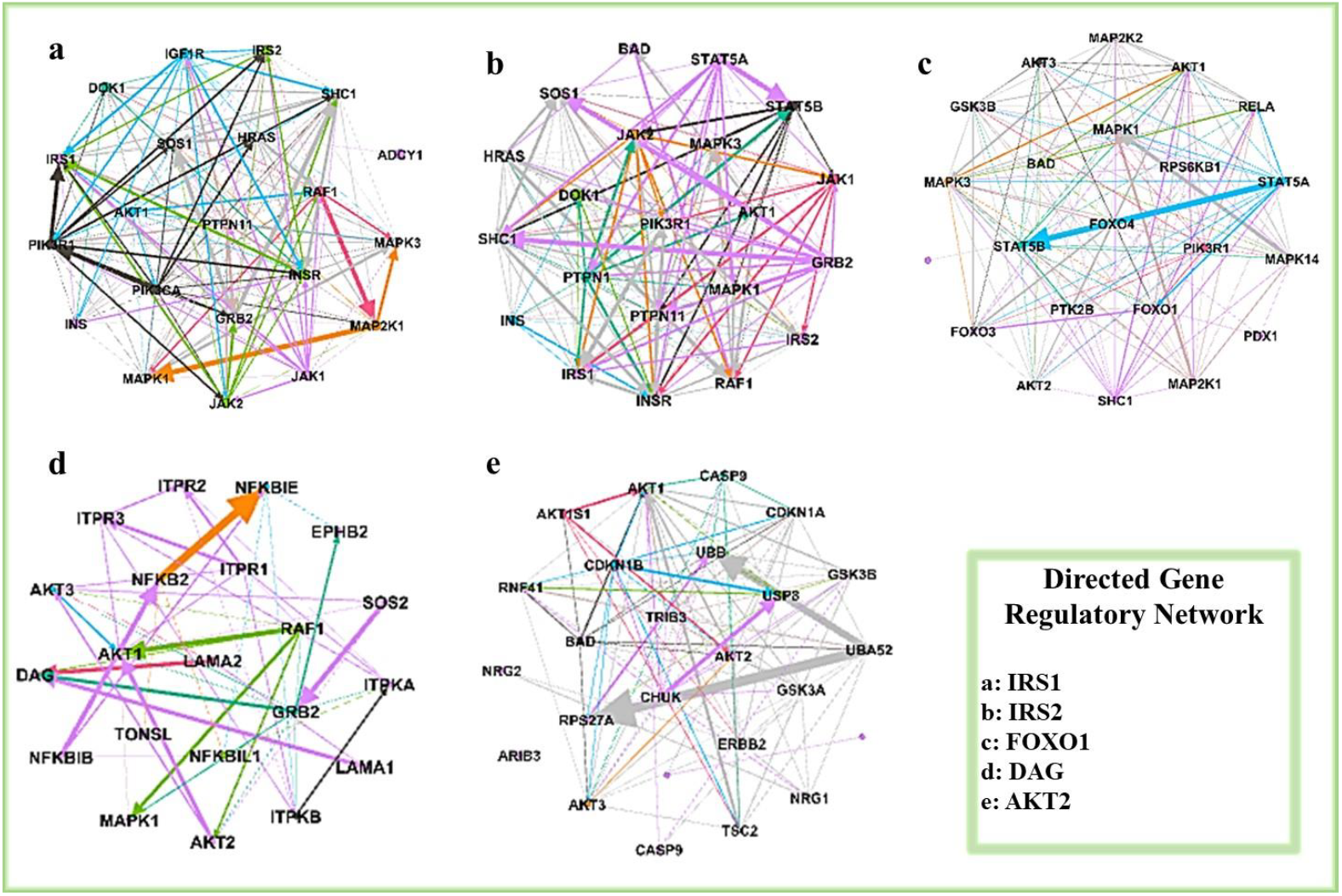
Directed gene regulatory networks of genes

Several genes were observed to be interacting with 5 key genes selected from published literature showing association with insulin resistance and metabolic diseases. Through Directed Gene Regulatory Network analysis and application of statistical test results, we were able to filter 18 Hub genes UBB, UBA52, NRG1, AKT3, NFKBIL1, AKT1, RELA, MAPK1, MAPK14, PTK2B, GSK3B, MAPK3, DOK1, SOS1, RAF1, SHC1, INSR and PIK3R1. These genes were filtered based on statistical test results values Closeness Centrality (CC) > 0.5); Harmonic closeness Centrality (HC) > 0.5); Eigen Centrality (EC) > 0.1); and Betweenness Centrality (BC) > 1) (Table 3).

**Table 3:** Identified Hub genes through gene regulatory network.

| S. No. | Genes | CC value | HC value | EC value | BC value | Hub |
| --- | --- | --- | --- | --- | --- | --- |
| 1 | UBB | 0.7 | 0.785714 | 0.310124 | 18.08333 | 0 |
| 2 | UBA52 | 0.714286 | 0.8 | 0.147258 | 22.75 | 0 |
| 3 | NRG1 | 1 | 1 | 0.284012 | 8 | 0 |
| 4 | AKT3 | 0.666667 | 0.75 | 0.356075 | 4 | 0.111747 |
| 5 | NFKBIL1 | 1 | 1 | 0.126848 | 2.5 | 0.075563 |
| 6 | AKT1 | 1 | 1 | 0.816136 | 7 | 0.045729 |
| 7 | RELA | 0.8 | 0.875 | 0.198619 | 11.12349 | 0.198842 |
| 8 | MAPK1 | 0.666667 | 0.791667 | 0.364812 | 20.10043 | 0.157825 |
| 9 | MAPK14 | 0.727273 | 0.8125 | 0.165843 | 16.45758 | 0.168895 |
| 10 | PTK2B | 0.727273 | 0.854167 | 0.166394 | 3.783009 | 0.194849 |
| 11 | GSK3B | 0.571429 | 0.760417 | 0.520458 | 18.7 | 0.164619 |
| 12 | MAPK3 | 0.833333 | 0.9 | 0.428249 | 2.490079 | 0.10664 |
| 13 | DOK1 | 1 | 1 | 0.392475 | 1.392857 | 0.111167 |
| 14 | SOS1 | 1 | 1 | 0.350811 | 4.123413 | 0.155442 |
| 15 | RAF1 | 0.833333 | 0.9 | 0.102 | 4.393687 | 0.215993 |
| 16 | SHC1 | 1 | 1 | 0.331766 | 18.94607 | 0.214087 |
| 17 | INSR | 0.615385 | 0.6875 | 0.68485 | 16.50043 | 0.083718 |
| 18 | PIK3R1 | 1 | 1 | 0.133489 | 2.946068 | 0.240457 |

CC (Closeness Centrality > 0.5); HC (Harmonic closeness Centrality > 0.5); EC (Eigen Centrality > 0.1); BC (Betweenness Centrality > 1).

### Functional annotation and pathway cross-talk network

The enrichment analysis method was used to identify over-represented classes of genes or proteins within a large group of samples, to reveal any existing associations with disease phenotypes. One approach to this was to use GO enrichment analysis, which can help to identify functionally enriched pathways of hub genes by examining their involvement in biological processes, molecular functions, and cellular components. Functional enrichment analysis is another approach, which applies statistical tests to match genes of interest with certain biological functions. By identifying these functional associations, enrichment analysis can help to shed light on the underlying mechanisms driving disease phenotypes and provide potential targets for therapeutic intervention. Every biological function has 50 gene ontology were shown in supplementary Table 4.

To identify functionally enriched pathways associated with hub genes, GO enrichment analysis was performed for biological process, molecular function, and cellular components. The enrichment analysis was validated using TOPPGENE and literature. Only statistically significant pathways (as determined by the FDR B&H test) and positive values greater than 1 were selected. These significant pathways and hub genes were then used to generate a network of functionally significant pathways with cross-talk between them (Figure 5).

**Figure 5:**
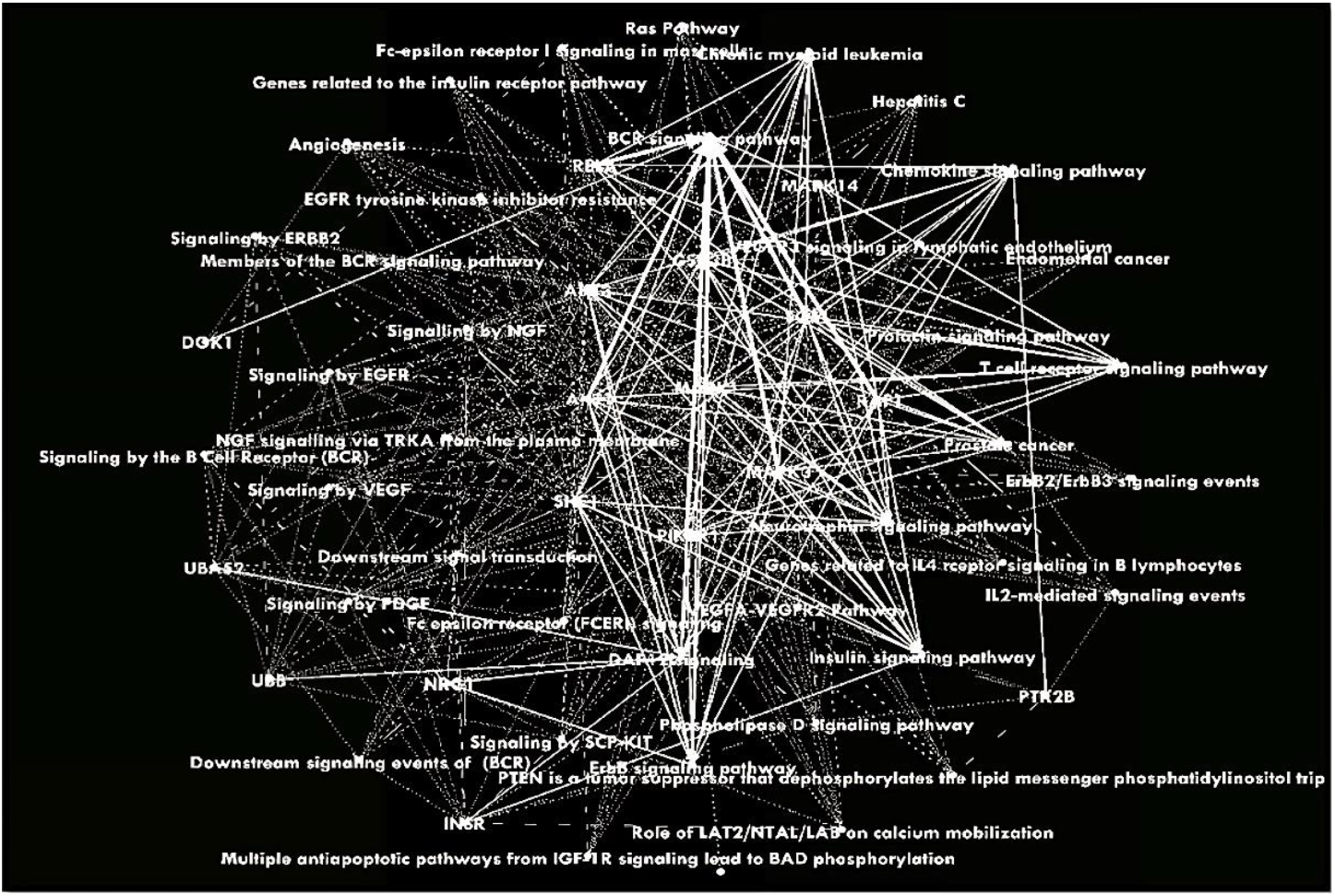
Pathway cross-talk generation of functionally enriched pathways of hub genes

### Validation of hub genes

An enrichment analysis was performed on a set of 18 genes, which were found to be enriched in functionally diverse pathways. Among these 18 genes, 8 hub genes were validated through literature and all of the hub genes were found to have a direct or indirect role in inflammation (Table 4).

**Table 4:** Validation of genes through literature how they play role in inflammation

| Sr.no | Gene | Role in insulin resistance | References |
| --- | --- | --- | --- |
| 1. | SHC1 | p66Shc is an adaptor protein that connects surface receptors to intracellular signaling pathways, influencing cellular response to nutrient availability. It is resistant to diabetes and obesity but inhibits glucose metabolism, promoting a shift towards anabolic metabolism. By counteracting insulin and mTOR signaling, p66Shc limits glucose uptake and metabolism, differentiating it from other Shc1 isoforms. | (29) |
| 2. | AKT1 | AKT1 is a key mediator of insulin, IGF1, and glucose responses and regulates muscle hypertrophy and atrophy. We examined the role of AKT1 variants in metabolic syndrome endophenotypes and found that insulin activated all Akt isoforms in lean muscles, while only Akt-1 was activated in obese muscles. Reduced IRS-1 expression and PI3K activity were observed in obese muscles, which are essential for insulin-induced Akt activation. | (12,30) |
| 3. | PIK3R1 | PIK3R1 regulatory subunits of PI3Ks are crucial for insulin signaling. Fibroblast studies of individuals with PIK3R1 mutations revealed severe insulin resistance in proximal and distal PI3K-dependent signaling. This sheds light on the genetic basis of severe insulin-resistance syndromes and highlights PIK3R1's role in normal development and diseases, including growth delay, ocular disorders, insulin resistance, diabetes, paucity of fat, and ovarian cysts. | (31) |
| 4. | GSK3B | GSK-3β is an enzyme that is suppressed by insulin. Elevated levels of GSK-3β result in insulin resistance and diabetes. Its role in beta cell development and function is poorly understood. GSK-3β deficiency in beta cells may control their growth through feedback inhibition of the insulin receptor/PI3K/Akt signaling pathway. | (18) |
| 5. | AKT3 | Resistin plays a role in insulin resistance by reducing insulin-stimulated glucose oxidation and decreasing IRS-1 and Akt1 function, leading to reduced glucose uptake and GLUT4 translocation. This may contribute to the pathophysiology of type 2 diabetes in obesity. | (32) |
| 6. | RELA | TRB1 acts as a transcriptional coactivator for RELA in adipocytes, promoting the production of proinflammatory cytokines. This dual role of TRB1 may contribute to the amplification of proinflammatory responses in white adipose tissue, which is often seen in sepsis, insulin resistance, and obesity-related type 2 diabetes. | (33) |
| 7. | MAPK1 | The role of p38 MAPK in regulating glucose transport in insulin-sensitive skeletal muscle, potentially through increased GLUT-4 activity, is not universally accepted. The specific role of p38 MAPK in regulating insulin-stimulated glucose transport in cultured myocytes is unclear. | (34,35) |
| 8. | INSR | Obesity is linked to insulin resistance and inflammation, and low-dose insulin infusion triggers a response in circulating human mononuclear cells (MNC). MNCs from obese individuals have reduced levels of phosphorylated insulin receptor beta subunit (p-INSR-beta), increased levels of pro-inflammatory mediators (IKBKB, SOCS, and PRKCB2) that are associated with p-INSR-beta, and reduced insulin sensitivity. | (36) |

## Discussion

The results presented describe a systematic approach to identifying core genes, interaction networks, hub genes, functional annotations, and pathway cross-talk networks associated with insulin resistance and metabolic diseases.

Insulin resistance is a condition characterized by decreased responsiveness of cells to insulin, leading to impaired glucose uptake and metabolism (37,38). Several genes have been implicated in the pathogenesis of insulin resistance and metabolic diseases, including IRS1, IRS2, AKT2, FOXO1, TNF-α, DAG, and IKK-β (31,32,39).

IRS1 and IRS2 are two critical genes that play a role in insulin signaling. These genes encode proteins that are involved in the activation of downstream signaling pathways that regulate glucose metabolism and lipid homeostasis (38). Both IRS1 and IRS2 are expressed in multiple tissues, including adipose tissue, liver, and skeletal muscle. In insulin-resistant states, these genes are often dysregulated, leading to impaired insulin signaling and glucose uptake (40,41).

AKT2 is another important gene involved in insulin signaling, and it plays a crucial role in glucose metabolism and lipid homeostasis. AKT2 is primarily expressed in insulin-responsive tissues such as skeletal muscle, adipose tissue, and liver. Defects in AKT2 have been linked to insulin resistance and type 2 diabetes (38,42).

FOXO1 is a transcription factor that regulates the expression of genes involved in glucose and lipid metabolism. FOXO1 is primarily expressed in liver and muscle tissue and is regulated by insulin signaling. In insulin-resistant states, FOXO1 activity is often dysregulated, leading to increased hepatic glucose production and impaired glucose uptake in muscle tissue (41,43)

TNF-α is a pro-inflammatory cytokine that plays a key role in the development of insulin resistance and metabolic diseases (44). TNF-α is primarily produced by adipose tissue and is involved in the regulation of adipocyte function and insulin signaling. Increased TNF-α levels have been linked to insulin resistance, and TNF-α inhibitors have shown promise in the treatment of metabolic diseases (40,45).

DAG is a second messenger molecule that is involved in insulin signaling. DAG activates protein kinase C (PKC), which in turn inhibits insulin signaling and impairs glucose uptake. In insulin-resistant states, DAG levels are often elevated, leading to impaired insulin signaling and glucose metabolism (43,46)

IKK-β is a kinase that is involved in the regulation of the NF-κB pathway, a key regulator of inflammation. IKK-β is primarily expressed in immune cells but is also present in other tissues, including adipose tissue and liver. In insulin-resistant states, IKK-β activity is often elevated, leading to increased inflammation and impaired insulin signaling (41,45).

Through our systematic bioinformatic approach we identified 18 hub genes (UBB, UBA52, NRG1, AKT3, NFKBIL1, AKT1, RELA, MAPK1, MAPK14, PTK2B, GSK3B, MAPK3, DOK1, SOS1, RAF1, SHC1, INSR, and PIK3R1) that play crucial roles in regulating the activity of other genes and pathways within the network. We were able to validate 8 genes SHC1, AKT1, PIK3R1, GSK3B, AKT3, RELA, MAPK1, and INSR of these 18 genes through published literature for playing a role in inflammation, either directly or indirectly that leads to pathogenesis of insulin resistance and metabolic diseases.

SHC1 is known to be involved in insulin signaling and has been shown to be a potential therapeutic target in insulin resistance (47). AKT1, on the other hand, has been implicated in regulating glucose homeostasis, lipid metabolism, and inflammation (48). PIK3R1 has also been found to play a role in insulin resistance by regulating glucose uptake and glycogen synthesis (47). GSK3B is a key regulator of insulin signaling and glucose metabolism, and its dysregulation has been linked to the development of insulin resistance and metabolic diseases (49). AKT3 has also been reported to play a role in insulin signaling and glucose metabolism (50). RELA, a subunit of the transcription factor NF-κB, has been implicated in inflammation and insulin resistance (51). MAPK1 has been shown to regulate glucose homeostasis and insulin sensitivity, while INSR has been identified as a key regulator of glucose uptake and insulin signaling (50).

Overall, these genes play crucial roles in regulating insulin signaling, glucose metabolism, and inflammation, and their dysregulation has been linked to the development of insulin resistance and metabolic diseases.

### Conclusion

The study aimed to investigate the role of insulin resistance in the onset of metabolic syndrome. To achieve this, the researchers identified key pathways and core genes involved in insulin resistance pathways and retrieved gene interaction data. They then identified and validated hub genes, which are genes that play a central role in the network of interactions between genes. We identified 18 hub genes that were involved in insulin resistance pathways. These genes were UBB, UBA52, NRG1, AKT3, NFKBIL1, AKT1, RELA, MAPK1, MAPK14, PTK2B, GSK3B, MAPK3, DOK1, SOS1, RAF1, SHC1, INSR and PIK3R1. To validate these hub genes, we looked for evidence in the literature that supported their involvement in insulin resistance or related pathways. we found that eight of the hub genes (SHC1, AKT1, PIK3R1, GSK3B, AKT3, RELA, MAPK1, and INSR) had been previously identified as playing a role in inflammation, either directly or indirectly. SHC1 is involved in insulin signaling and glucose metabolism. AKT1, a key player in the insulin signaling pathway, regulates various cellular processes crucial for glucose metabolism and cell survival. PIK3R1, a regulatory subunit of PI3K, contributes to AKT activation and downstream signaling, influencing glucose metabolism. GSK3B is essential for proper insulin signaling and glycogen synthesis, and its dysregulation is associated with insulin resistance. AKT3, another member of the AKT family, also plays a role in insulin signaling and glucose metabolism. RELA, a subunit of NF-κB, contributes to the regulation of inflammation, linking chronic inflammation to metabolic disorders. MAPK1, part of the MAPK signaling pathway, is implicated in insulin resistance and inflammation. Lastly, INSR, the insulin receptor, is central to insulin signaling, and dysregulation of its function is associated with insulin resistance and metabolic disorders. These genes collectively participate in complex networks that influence inflammation and metabolic processes, with their dysregulation contributing to the development and progression of metabolic diseases.

These findings suggest that inflammation plays a crucial role in the pathogenesis of insulin resistance, leading to the development of metabolic diseases or metabolic syndrome. The identified hub genes may serve as potential targets for future therapeutic interventions to prevent or treat metabolic syndrome. Insulin resistance is the main component of metabolic syndrome that paly role in inflammation.

### Future Perspective

The identification of hub genes and key pathways involved in insulin resistance is an important step towards understanding the underlying mechanisms of metabolic syndrome. However, further studies are necessary to validate the findings and determine how the identified genes and pathways contribute to the development of insulin resistance in different populations and individuals.

One possible future direction for this project is to conduct a genetic analysis of the identified hub genes in different populations and individuals. This can be done using PCR (polymerase chain reaction) to amplify and detect specific DNA sequences associated with the hub genes. By analyzing the genetic variation in these genes, researchers can determine which alleles are present and how they contribute to insulin resistance and metabolic syndrome.

This type of genetic analysis can provide valuable information for developing personalized interventions and treatments for metabolic syndrome. For example, individuals with certain alleles of the identified hub genes may benefit more from specific dietary or exercise interventions or may require different medication regimens. Additionally, understanding the genetic basis of insulin resistance can help identify new drug targets and improve the development of novel therapies for metabolic syndrome.

## Conflict of Interest

There are no conflicts of interest related to this research.

## Notes

### Competing Interest Statement

The authors have declared no competing interest.

